# Cardiac and respiratory muscle responses to dietary N-acetylcysteine in rats consuming a high-saturated fat, high-sucrose diet

**DOI:** 10.1101/2021.06.02.446720

**Authors:** Rachel C. Kelley, Derek R. Muscato, Dongwoo Hahn, Demetra D. Christou, Leonardo F. Ferreira

**Affiliations:** Department of Applied Physiology and Kinesiology, University of Florida, Gainesville, FL

## Abstract

**BACKGROUND:** Exertional dyspnea is a significant clinical concern in individuals with overweight or obesity. The pathophysiology of dyspnea is multifactorial and complex. Previous data suggest that diaphragm and cardiac abnormalities should be considered as likely contributors to obesity-related exertional dyspnea. Additionally, oxidative stress is a causative factor in the general etiology of obesity as well as skeletal and cardiac muscle pathology. Thus, this preclinical study aimed to define diaphragm and cardiac morphological and functional alterations following an obesogenic diet in rats and the therapeutic potential of an antioxidant supplement, N-acetylcysteine (NAC).

**METHODS:** Male Wistar rats (∼7 weeks old) consumed ad libitum either lean (20% protein, 70% carbohydrate, 10% fat) or high-saturated fat, high-sucrose (HFHS, 20% protein, 35% carbohydrate, 45% fat) diets for ∼22 weeks. Rats receiving HFHS diet were randomized to drink control water or water with NAC (2 mg/ml) for the last eight weeks of the dietary intervention: Lean, HFHS, and HFHS+NAC (n = 8 per group). We evaluated diaphragm bundles (in vitro function and histology) and hearts (weights and echocardiography) for all groups.

**RESULTS:** Final body weights of HFHS rats, but not HFHS+NAC rats, were significantly higher than Lean controls. Neither HFHS diet nor NAC supplementation affected diaphragm specific force (N/cm^2^), peak power (W/kg), or morphology. In cardiac muscle, right and left ventricle weights (normalized to tibia length) of HFHS rats were greater than those of Lean controls and HFHS+NAC rats. Cardiac functional abnormalities were also present in HFHS rats, with left ventricular fractional shortening (%) and posterior wall maximal shortening velocity (cm/s) increasing compared to Lean controls, but HFHS+NAC rats did not demonstrate these markers of hypercontractility. HFHS rats showed an elevated deceleration rate of early transmitral diastolic velocity (E/DT) consistent with diastolic dysfunction, but NAC eliminated this effect.

**CONCLUSION:** Our data suggest that an HFHS diet does not compromise diaphragm muscle morphology or in vitro function, suggesting other possible contributors to breathing abnormalities in obesity (e.g., neuromuscular transmission abnormalities). However, an HFHS diet resulted in cardiac hypertrophy, hypercontractility, and diastolic dysfunction. Supplementation with NAC did not affect diaphragm morphology or function but attenuated cardiac abnormalities in the HFHS diet. Our findings support future studies testing NAC supplementation in clinical trials of humans with obesity.

## Introduction

Dyspnea is a common clinical concern for individuals with overweight or obesity.^1^ This discomfort and breathlessness typically manifest during physical activity, thus compromising an individual’s exercise capacity.^2-4^ Exercise, regardless of weight loss, improves morbidity and mortality in individuals with obesity, so there is an urgent clinical need to address barriers to physical activity.^5,6^ Obesity-related dyspnea can also impair lower intensity activities of daily living.^7^ Indeed, 80% of individuals with obesity reported shortness of breath after climbing two flights of stairs compared to 16% of normal-weight participants.^8^ Exercise intolerance, difficulties in daily tasks, and even the anticipation of dyspnea inevitably decrease an individual’s quality of life.^9^

The pathophysiology of dyspnea is multifactorial and complex,^10^ but two factors worthy of investigation in obesity specifically include respiratory muscle weakness and cardiac dysfunction. Elevated body weight increases work and oxygen cost of breathing in obesity, likely through a diminished chest wall or lung compliance.^2-4,11^ Increased work of breathing would require respiratory muscles to expend more energy, fatigue faster, and send afferent feedback that would contribute to perceived shortness of breath.^2^ Any abnormalities in respiratory muscles (i.e., atrophy or contractile dysfunction) would further exacerbate dyspnea during exertion.^12,13^ Researchers have reported impaired maximal inspiratory pressure, a clinical marker of inspiratory (diaphragm) muscle strength, in individuals with obesity.^14-19^ This finding suggests that diaphragm weakness may be a relevant therapeutic target for attenuating dyspnea and improving exercise capacity in this patient population. Preliminary data show improvements in exercise performance and tolerance when individuals with obesity undergo respiratory muscle unloading^20^ or training.^21-23^ Additionally, animal data have supported the notion of diaphragm abnormalities in obesity. Obese rodents have demonstrated diaphragm weakness,^24-27^ slow fiber type shifts,^24,28^ increased fibrosis,^29^ and/or atrophy of fibers.^29^

Cardiac abnormalities are also important drivers of obesity-related dyspnea. Individuals with impaired myocardial contractility often develop pulmonary congestion, which increases the work of breathing and also stimulates sensory neurons that reflexively augment ventilation and lead to perceived breathlessness.^10,30^ Importantly, cardiac remodeling occurs in persons with obesity and contributes to their exacerbated risk for cardiovascular diseases, such as heart failure.^31,32^ Individuals with obesity commonly exhibit abnormalities in cardiac morphology, including fatty infiltration, fibrosis, and concentric cardiac hypertrophy.^32-35^ These morphological changes are associated with altered systolic and diastolic function.^32,33^ Interestingly, it appears that obesity contributes to this cardiac pathology independent of other cardiometabolic syndrome risk factors, such as high blood pressure and type 2 diabetes.^35^

Oxidative stress is a likely mechanism contributing to this diaphragm and cardiac dysfunction in obesity. Redox imbalance plays a major role in the etiology of obesity and metabolic syndrome.^36-38^ Furthermore, obesogenic diets have been shown to alter markers of reactive oxygen species emission and antioxidant system capacity in skeletal muscle and myocardium in rodents.^39-47^ Clinical trials exploring antioxidant interventions, like supplementation with vitamins C and E, in obesity have broadly reported null or negative results.^48^ Selection of vitamins as antioxidants may not be ideal because they scavenge oxidants in a stoichiometric manner and therefore cannot lead to long-term prevention of reactive oxygen species formation.^48^ An alternative antioxidant supplement is N-acetylcysteine (NAC). Unlike antioxidant vitamins, NAC contributes to the synthesis of glutathione, the main intracellular antioxidant. Thus, NAC can provide a chronic defense against oxidative stress. Furthermore, enzymes involved in glutathione synthesis are regulated by cellular redox state.^49^ This regulation means that NAC therapy may be less prone to inducing reductive stress than other antioxidant supplements.

Following this literature and rationale, we hypothesized that a high-saturated fat, high-sucrose (HFHS) diet in rats would cause diaphragm muscle and cardiac abnormalities. Macronutrient profiles of obesogenic diets have not been consistent across previous rodent studies. We sought to utilize a diet with a clinically relevant composition similar to a processed ‘Western’ style diet commonly consumed in the United States.^50,51^ Based on the role of oxidants on cardiac and diaphragm muscle dysfunction and systemic pathophysiology of HFHS diet, we predicted that N-acetylcysteine (NAC), which has antioxidant properties, would have therapeutic effects.

## Methods

### Animals and diet

Adult male Wistar rats (initially 7-9 weeks old) were used in this study. Rats were housed at the University of Florida under 12h:12h light-dark cycle and had access to their assigned rodent diet and water ad libitum. Rats were pair-housed for approximately 6 weeks but were then separated for most of the study for accurate measurement of individual food and water intake. All animal studies were performed with approval from the University of Florida Institutional Animal Care and Use Committee (IACUC 201709714).

All rats underwent a 1-week acclimation period before being randomly allocated to a semi-purified, irradiated diet. Rats continued on the allocated diet for 20 to 24 weeks. Lean control rats (Lean, n = 8) had ad libitum access to a lean diet (3.85 kcal/g; % carbohydrate:fat:protein = 70:10:20; Research Diets, #D12450K). The HFHS group (n = 16) had ad libitum access to an obesogenic diet (4.73 kcal/g; % carbohydrate:fat:protein = 35:45:20, Research Diets, #D12451). Lean diet fat sources were soybean oil and lard (5:4 grams), and carbohydrate sources were corn starch and maltodextrin (11:3 grams). Notably, the lean diet did not have sucrose. HFHS diet fat sources were soybean oil and lard (25:178 grams), and carbohydrate sources were sucrose, corn starch, and maltodextrin (177:73:100 grams). We weighed the rats, the amount of food provided, and the amount of food remaining once per week. Apparent daily diet volume (g) and energy (kcal) consumption were calculated from these weekly measurements.

### N-acetylcysteine treatment

Approximately 8 weeks before terminal experiments, a subset of HFHS rats began receiving N-acetylcysteine (NAC) in drinking water. These rats (HFHS+NAC, n = 8) were randomly allocated to this treatment designation at the beginning of the study. Stock aliquots of NAC (100 mg/ml) in deionized water were prepared and frozen at - 20°C. Every day, frozen aliquots were removed, thawed, and mixed with reverse osmosed water for a final concentration of 2 mg/ml in drinking water. We prepared ‘fresh’ NAC solutions daily for drinking to minimize the confounding effects of auto-oxidation on the antioxidant properties of NAC. Rats typically consume ∼30 to 40 ml of water daily.^52^ Therefore, this concentration of NAC would result in ∼75 mg NAC/day, which is a dose that has previously been shown to resolve superoxide overproduction and contractile dysfunction in cardiomyocytes isolated from rats that have undergone myocardial infarction surgery.^52^

### Glucose tolerance testing

Rats underwent glucose tolerance testing approximately 1 to 2 weeks before terminal experiments. Glucose tolerance tests were performed by intraperitoneal injection of glucose (2 g/kg, 20% w/v D-glucose, in 0.9% w/v saline) following a 6-hour fast. Blood samples were taken from the tail vein immediately before (time 0 min), and 15 min-, 30 min-, 60 min-, 90 min-, and 120-min post-glucose bolus. Blood glucose concentration was determined by a glucometer (Abbott, FreeStyle Lite). All fasts prior to glucose tolerance testing began at 0800 h, and fasting blood glucose was measured at 1400 h (time 0 min).

### Echocardiography

Rats underwent cardiac ultrasound for evaluation of left ventricle systolic and diastolic function. Animals were maintained at 1.5% to 2.5% isoflurane anesthesia, and ECG electrodes were placed on the limbs to monitor heart rate. The level of anesthesia was adjusted as needed to maintain a heart rate of ∼ 400 bpm. Two-dimensional ultrasound images were obtained using M-mode and pulse wave Doppler imaging with a 7.5 MHz sector transducer (Aplio XV, Toshiba America Medical Systems, Tustin, CA, USA). Image J software was used to analyze the images. M-mode parasternal short-axis images were used to quantify left ventricular internal diameters during diastole (LVIDd) and systole (LVIDs). Pulse wave Doppler 4-chamber apical images at the mitral valve level were used to quantify E wave velocity (E) and E wave deceleration time (DT). Notably, the rodents’ rapid heart rates often cause E and A waves to fuse unless anesthesia is increased to lower heart rate substantially. Therefore, only the fused peak “E” velocity was measured in this study as previously advised in recommendations for echocardiographic examination in rodents.^53^ Fractional shortening (FS%) was defined as (LVIDd-LVIDs)/LVIDd x 100, and E wave deceleration rate was defined as E/DT. Importantly, E wave deceleration rate has been shown to correlate positively with restrictive filling and pulmonary congestion.^54^ All ultrasound imaging was conducted at approximately 1700h to 2100h.

### Tissue Collection

All researchers were blinded to group assignment for tissue collection and accompanying assays and statistical tests. Rats were fasted prior to terminal experiments and randomly allocated for sacrifice at 0830h or 1300h. We anaesthetized rats using isoflurane and performed a laparotomy and thoracotomy to collect tissue samples. All dissected tissues were placed in ice-cold Krebs Ringer solution (in mM: 137 NaCl, 5 KCl, 1 MgSO_4_, 1 NaH_2_PO_4_, 24 NaHCO_3_, and 2 CaCl_2_). The diaphragm was excised, and then the heart was removed and dissected to separate right ventricle and left ventricle + septum. The costal diaphragm was processed and allocated for contractile and morphological assays.

### Diaphragm muscle contractile function

We dissected a bundle of the costal diaphragm with rib and central tendon remaining for attachment to a muscle mechanics apparatus (Aurora Scientific, 300C L-R model) and analysis of contractile properties as previously described.^55,56^ The bundle was kept in Krebs Ringer solution gassed with a mixture of 95% O_2_ and 5% CO_2_ throughout the procedure. We determined the passive length-tension relationship and found optimal length for isometric contraction using a process previously reported in mice.^57^ Briefly, the bundle was slowly stretched by moving the lever arm until passive force reached ∼25 mN. The lever arm position was recorded, and the muscle supramaximally stimulated repeatedly (1 Hz, 600 mA, 0.25 ms pulse) at one-minute intervals, with the bundle being progressively shortened by 0.3 mm and force allowed to reach a steady-state before each stimulation. Peak and baseline forces were recorded, and the muscle was placed at the length that elicited the highest active (peak - baseline) force (optimal length, l_o_). The preparation was then warmed from room temperature to 37°C, allowing 10 minutes for thermo-equilibration. Using the same current and pulse as stated above, the isometric force was measured at the following stimulation frequencies in order: 120 Hz (maximal), 1 Hz (twitch), and 40 Hz (submaximal). This abbreviated stimulation protocol was selected to maintain preparation stability prior to isotonic contraction and prevent rundown of force, which may occur during full force-frequency evaluation of diaphragm bundles. Five minutes after these contractions, the maximal tetanic force (P_o_) produced by each muscle was used as a reference for an isotonic release protocol as described previously.^57^ In the isotonic release experiment, the diaphragm underwent maximal isometric contraction for 300 ms and then was allowed to shorten by reducing the load to 30% to 35% P_o_, which is the load that approximately elicits peak power.^58,59^ We normalized force by the diaphragm bundle cross-sectional area (CSA, N/cm^2^). To estimate the diaphragm bundle CSA, we divided bundle weight (g) by bundle length (cm) multiplied by the muscle specific density (1.056 g/cm^3^).^60^ To calculate diaphragm peak power (in W/kg), we multiplied force generated during shortening (N/kg) x velocity (m/s).

### Diaphragm fiber cross-sectional area and fibrosis

On the day of terminal experiments, a bundle of left costal diaphragm allotted to immunohistochemistry and histology assays was embedded in Tissue-Tek OCT freezing medium, frozen in liquid-nitrogen-cooled isopentane, and stored at -80°C until later processing. Diaphragm bundles were processed for immunohistochemical analyses of fiber cross-sectional area and myosin heavy chain isoforms as described previously.^57,61^ Bundles embedded in Tissue-Tek OCT freezing medium were sliced into sections of 10 μm thickness at approximately -20ºC using a cryostat (Leica, CM 3050S model). Sections were incubated in 1:200 wheat germ agglutinin (WGA) Texas Red (Molecular Probes) for 1 hour at room temperature, washed in PBS 3 x 5 min, permeabilized with 0.5% Triton X-100 solution for 5 min, washed in PBS 1 x 5 min, and incubated in primary antibodies in a humid chamber for 90 min. Primary antibodies were myosin heavy chain (MyHC) type I (1:15) and MyHC type IIa (1:50). The MyHC I (A4.840) and MyHC IIa (SC-71) antibodies were developed by Stanford University and University of Padova, respectively, and obtained from the Developmental Studies Hybridoma Bank, created by the NICHD of the NIH and maintained at The University of Iowa, Department of Biology, Iowa City, IA 52242. After the primary antibody incubation, sections were washed in PBS 3 x 5 min and exposed to fluorescently conjugated secondary antibodies (60 min; Goat x Mouse IgM Alexa 350 and Goat x Mouse IgG Alexa 488, Invitrogen). Sections were then washed in PBS 3 x 5 min, allowed to dry, and imaged. We acquired and merged images using an inverted fluorescence microscope (Axio Observer, 10x objective lens) connected to a monochrome camera (Axio MRm, 1x c-mount, 2/3” sensor) and Zen Pro software (Carl Zeiss Microscopy). We used semi-automatic muscle analysis using segmentation of histology (SMASH) code, run in MATLAB software, to quantify fiber type distribution and fiber cross-sectional area for multiple images per rat.^62^ We analyzed 269-737 diaphragm fibers per rat.

We used a commercial kit (Newcomer’s Supply, #9179B) for Masson’s trichrome staining of diaphragm sections following the kit’s instructions with some modifications. Briefly, 10 μm-thick cross-sections were incubated in 4% paraformaldehyde at room temperature (15 min). After rinsing with distilled water, slides were then left in Bouin’s fluid overnight at room temperature. Slides were then washed with tap water followed by a distilled water rinse and then completed staining with the following solutions: hematoxylin (10 min), Biebrich Scarlet Fuchsin Satin (5 min), phosphomolybdic-Phosphotungstic acid (10 min), Anilin Blue (5 min), and 0.5% acetic acid (3 min). Rinses with water followed these incubations as per kit instructions. Slides were dehydrated with ethanol and cleared with xylene rinses before cover-slipping in mounting medium (Permount, Fisher). We acquired and merged images using an inverted fluorescence microscope (Axio Observer, 10x objective lens) connected to a color camera (Axio ERc 5s, 1x c-mount, 1/2.5” sensor) and Zen Pro software (Carl Zeiss Microscopy). Images were analyzed using Image J software (National Institutes of Health) for percent fibrosis.

### Statistics

We used One-Way ANOVA (Prism 6, GraphPad Software Inc., La Jolla, CA) for group comparisons. When One-Way ANOVA results were statistically significant, Lean and HFHS+NAC groups were compared to the HFHS as the fixed group using Dunnett’s post hoc tests. We thus tested the pathological potential of the HFHS diet (i.e., Lean vs. HFHS) and the therapeutic effects of NAC (i.e., HFHS vs. HFHS+NAC). Our Dunnett’s post hoc testing method did not analyze the Lean vs. HFHS+NAC comparison, and by decreasing the number of comparisons, we were able to increase our statistical power. We also used a paired Student’s t-test to evaluate differences in diastolic function before (Pre) and after (Post) NAC treatment. Parametric and non-parametric tests were used based on results from normality (Shapiro-Wilk) and equal variance (Brown-Forsythe) tests (Sigma Plot v.13, Systat). All data are shown as mean ± SD or scatter plots with bars indicating group means. In general, we used the conventional *p*-value of less than 0.05 to declare statistical significance, but wherever feasible, we report exact *p*-values. Importantly, we have taken into consideration recent recommendations for our data interpretation.^63^

## Results

### Animal Characteristics

Rats consuming the high-saturated fat, high-sucrose (HFHS) had ∼16% ± 12% higher terminal body weights than rats fed a lean control diet (Figure 1 A-B). This difference in body weight remained when normalized to tibia length (TL), a marker of body size, suggesting increases in fat stores. Notably, rats on the HFHS diet who received N-acetylcysteine (NAC) supplementation weighed less than those on the HFHS diet alone (Figure 1 B). There were no differences in energy intake across groups, but the HFHS+NAC rats consumed less water than those on the HFHS diet alone (Figure 1 E-F).

**Figure 1.**
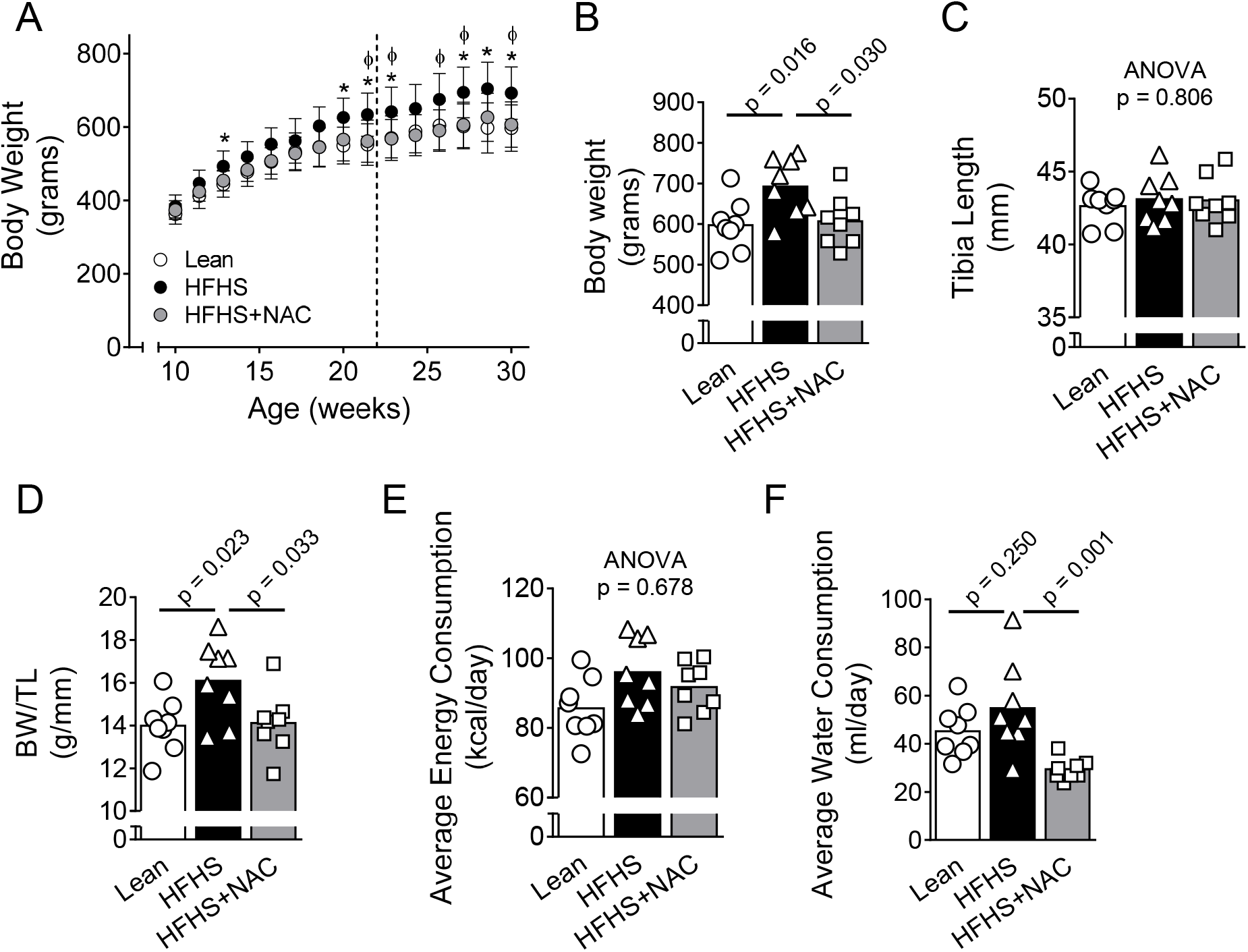
Body weight, energy intake, and water consumption. Lean = lean control diet, HFHS = high-saturated fat, high sucrose (HFHS) diet, N-acetylcysteine = NAC in drinking water. (A) Weight gain with age. The dotted line indicates the start of NAC treatment, which was ∼14 weeks after start of assigned diets. (B) Terminal body weights. (C) Terminal tibia length as a marker of body size. (D) BW/TL ratio as a marker of adiposity. (E) Average apparent daily energy intake. (F) Average apparent daily water consumption. Data are mean ± SD in panel A. In panels B-D, data are shown as scatter plots and mean bars. In panels E-F, data are shown as mean scatter plot data per animal, and bars are group averages of these means. Comparisons among 3 groups were conducted using One-Way ANOVA. Individual p values for post hoc tests (Dunnett) are shown when feasible. * = p<0.05 Lean vs HFHS, Φ = p <0.05 HFHS vs. HFHS+NAC. BW = body weight, TL = tibia length

Glucose handling measured with our protocol was not different among groups (Figure 2). Fasting blood glucose levels were within the normal range for all rats (Figure 2 B). However, HFHS rats had statistically higher fasting blood glucose readings than Lean controls (in mg/dl: Lean 63 ± 4.5, HFHS 79.5 ± 13, p = 0.003).

**Figure 2.**
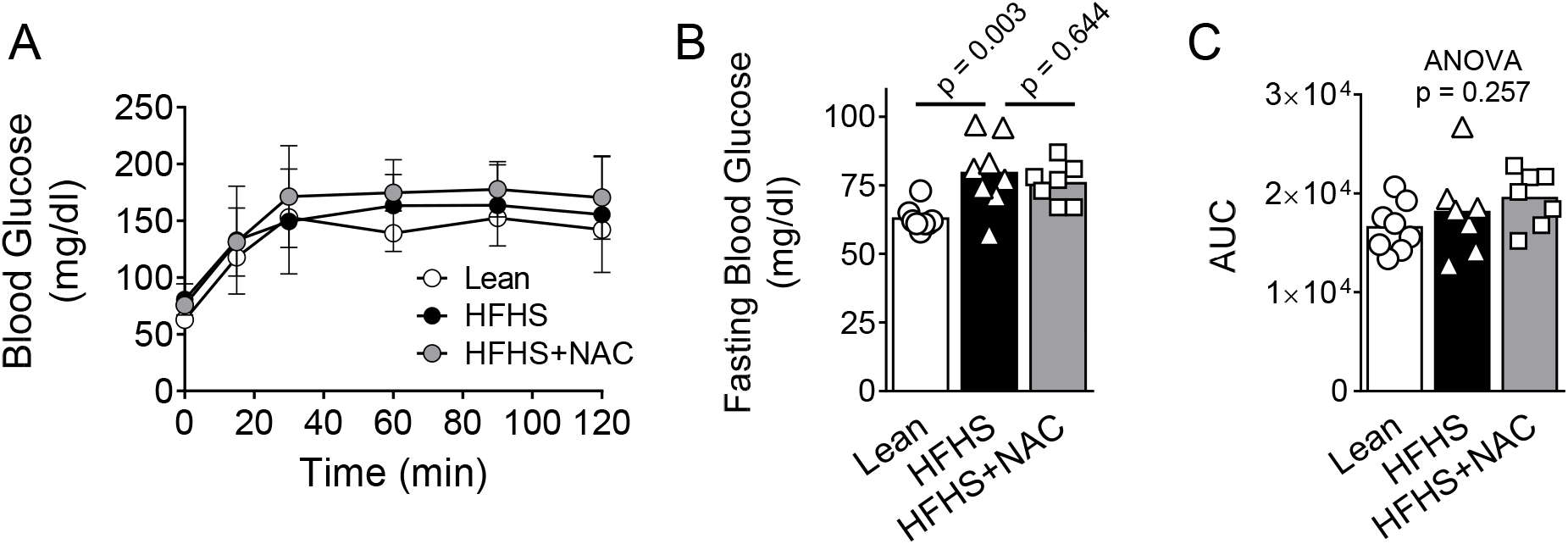
Glucose handling. (A) Glucose tolerance test. (B) Fasting blood glucose (6-hour fast). (C) Glucose tolerance test area under the curve (AUC) calculations. Data are mean ± SD in panel A. In panels B-C, data are shown as scatter plot and mean bars. Comparisons among 3 groups were conducted using One-Way ANOVA. Individual p values for post hoc tests (Dunnett) are shown when feasible.

### Diaphragm Contractile Function, Fiber Typing, Passive Mechanics, and Fibrosis

We did not observe differences in diaphragm isometric force production at twitch, submaximal, or maximal direct field stimulation (Figure 3 A-C). Twitch time-to-peak tension (TPT, in ms: Lean 19.5 ± 0.5, HFHS 20.9 ± 3.7, HFHS+NAC 19 ± 2.0, p = 0.32) and 1/2 relaxation time (1/2 RT, in ms: Lean 18.2 ± 2.0, HFHS 19.2 ± 3.7, HFHS+NAC 18.6 ± 4.9, p = 0.87) were unaltered. There were also no differences in maximal rate of contraction (in N/cm^2^•ms^-1^: Lean 0.82 ± 0.12, HFHS 0.74 ± 0.14, HFHS+NAC 0.74 ± 0.11, p = 0.37) nor maximal rate of relaxation (in N/cm^2^•ms^-1^: Lean 1.38 ± 0.18, HFHS 1.24 ± 0.19, HFHS+NAC 1.20 ± 0.19, p = 0.16) during tetanic contraction (120 Hz). When force was clamped at 30% to 35% of maximum and the bundle then allowed to shorten, we similarly found no statistically significant difference in diaphragm peak power (Figure 3 D). In parallel to the lack of these functional changes, there were no morphological differences in diaphragm muscle fiber size (Figure 4). Additionally, myosin heavy chain isoform distribution quantified using immunohistochemistry analyses did not differ among groups (%Type I: Lean 33 ± 5, HFHS 33 ± 7, HFHS+NAC 32 ± 5, p = 0.96; %Type IIa: Lean 35 ± 6, HFHS 35 ± 8, HFHS+NAC 36 ± 7, p = 0.98; %Type IIb/x: Lean 32 ± 6, HFHS 32 ± 9, HFHS+NAC 32 ± 4, p = 0.99).

**Figure 3.**
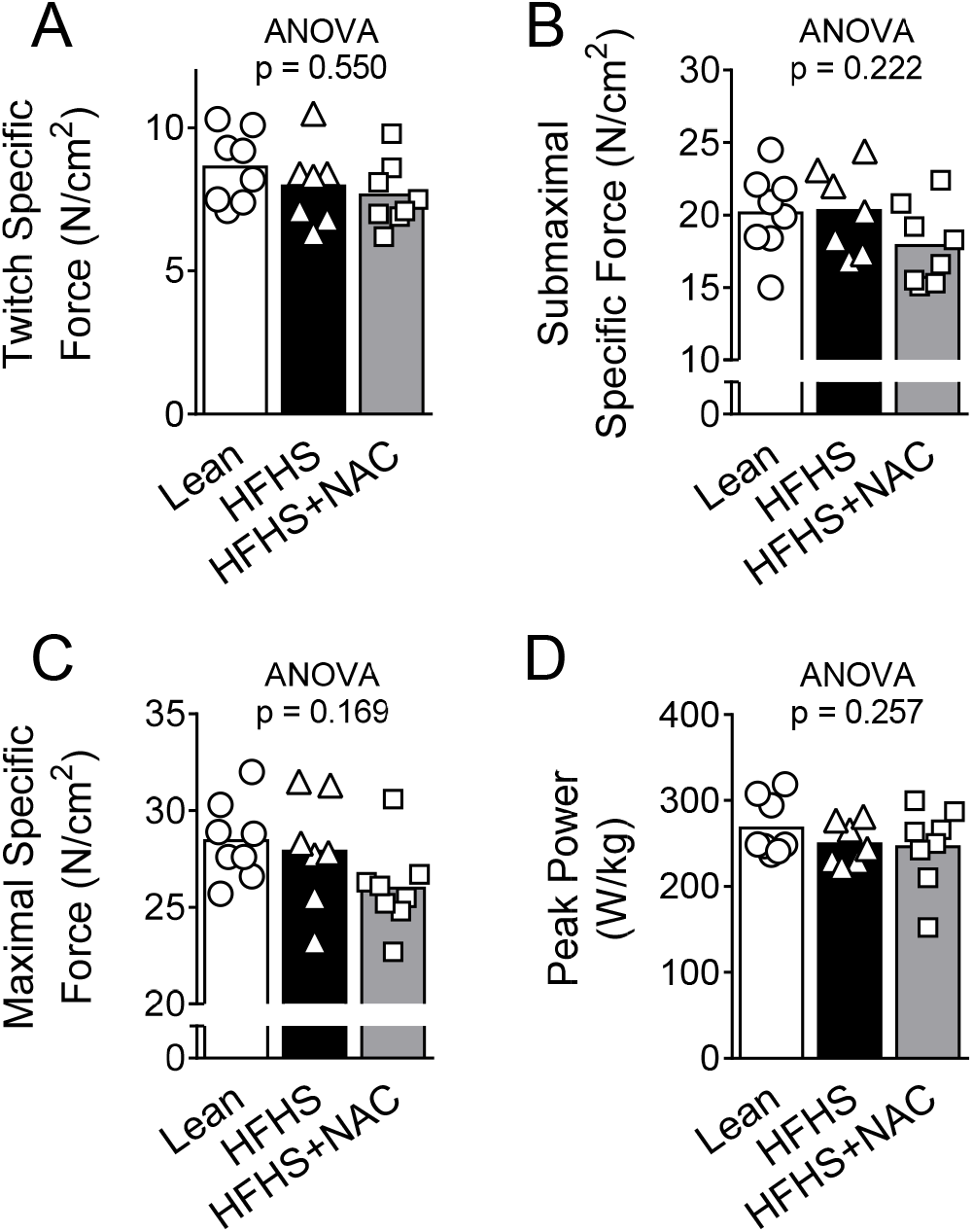
Diaphragm contractile function. (A) Twitch (1 Hz), (B) Submaximal (40 Hz), and (C) maximal (120 Hz) isometric specific force of diaphragm bundle. (D) Power of diaphragm bundle using a clamped load of 30%-35% maximal force. Comparisons among 3 groups were conducted using One-Way ANOVA. No comparisons surpassed the threshold for statistical significance (P < 0.05).

**Figure 4.**
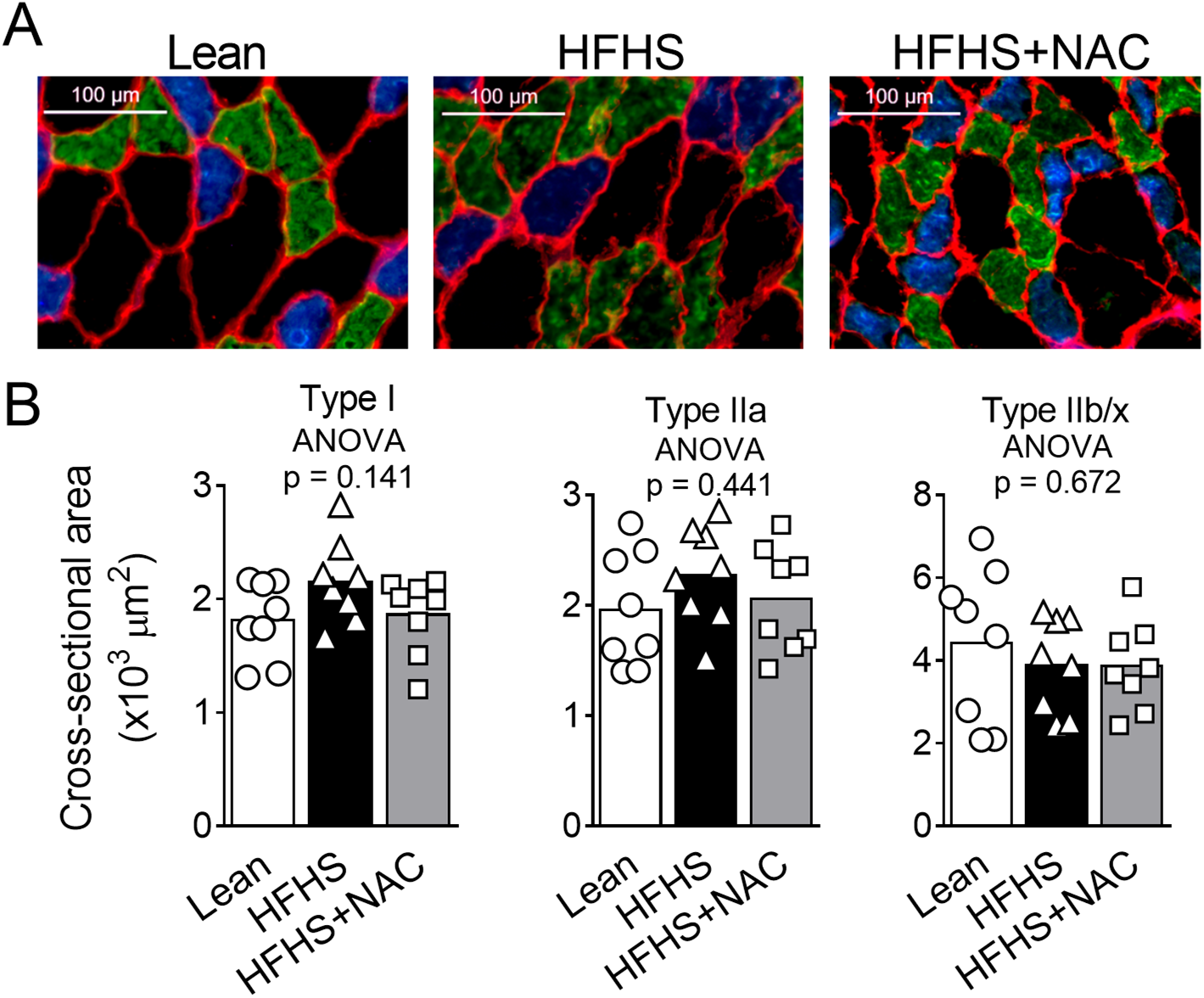
Diaphragm fiber cross-sectional area. Diaphragm fiber cross-sectional area and myosin heavy chain isoform distribution. (A) Sample images of diaphragm immunohistochemistry for fiber typing and cross-sectional area. Colors represent specific MHC isoforms (blue = type I, green = type IIa, black = type IIb/x). (B) Cross-sectional area for each fiber type in the diaphragm. Immunohistochemistry data are shown as scatter plots of the average value of individual fibers measured for each animal and bars representing group means. Comparisons among 3 groups were conducted using One-Way ANOVA. No comparisons surpassed the threshold for statistical significance (P < 0.05).

Masson’s Trichrome staining revealed no difference in fibrotic tissue content among groups (Figure 5 A-B). Relatedly, evaluation of diaphragm passive mechanical properties demonstrated that diaphragm bundle stiffness did not differ among groups (Figure 5 C-D).

**Figure 5.**
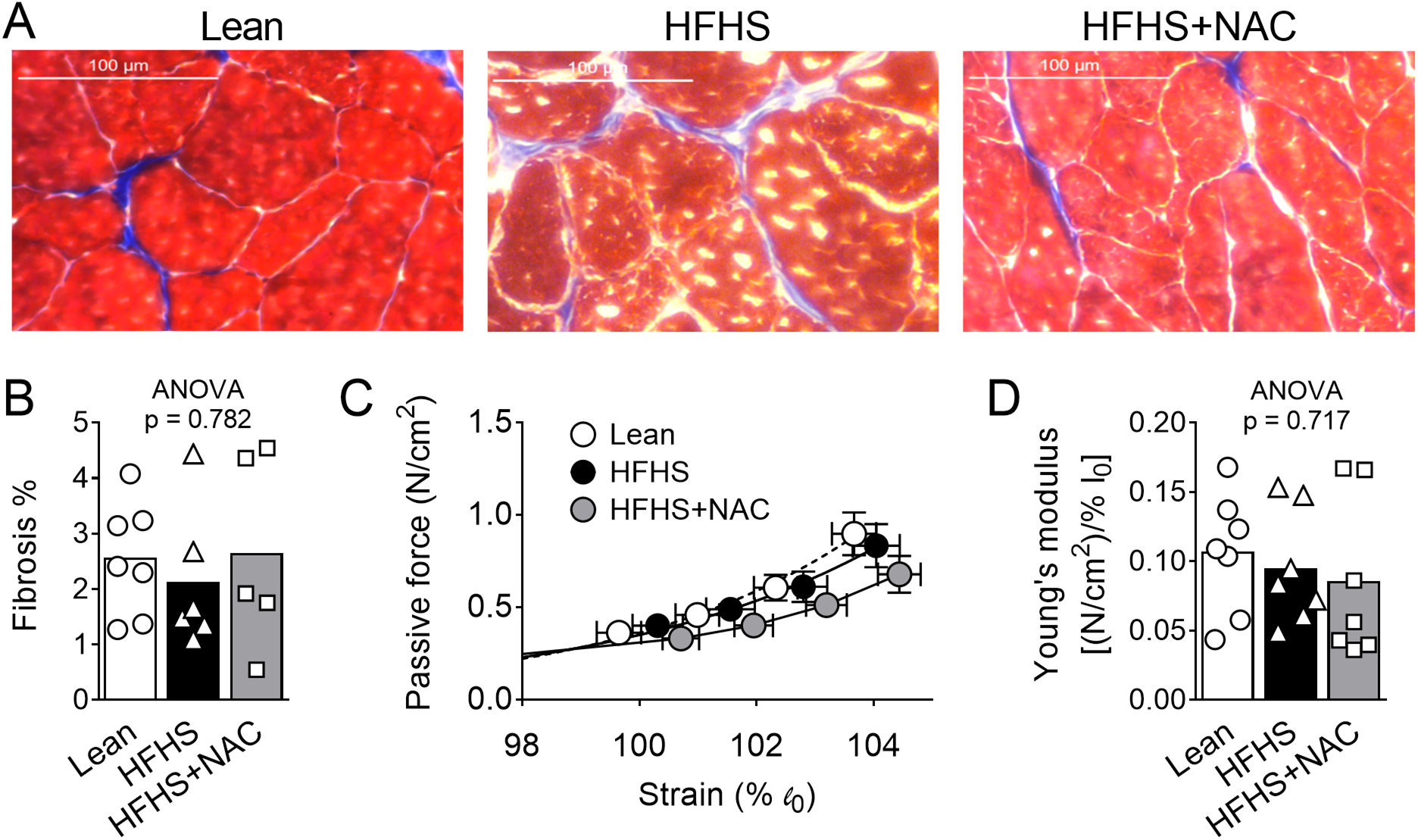
Diaphragm passive tension, stiffness, and fibrosis. (A) Sample images of diaphragm Masson’s Trichrome stain for quantification of fibrotic tissue. (B) Quantification of % fibrosis in diaphragm shown as scatter plots of the average value for each animal and bars representing group means. (C) Relationship between passive tension (N/cm^2^) and strain (normalized to optimal length) in diaphragm bundles. (D) Young’s elastic modulus of diaphragm bundles was calculated as the change in passive tension normalized to strain (∼3% from optimal length). Data are shown as scatter plots and mean bars. Comparisons among 3 groups were conducted using One-Way ANOVA. No comparisons surpassed the threshold for statistical significance (P < 0.05).

### Cardiac Size and Function

Rats underwent echocardiography evaluation of left ventricular function after ∼14 weeks on the assigned diet before any NAC treatment. Table 1 shows that there were no differences in left ventricular diameters or indices of left ventricular systolic (i.e., fractional shortening and posterior wall shortening velocity) and diastolic (i.e., E wave, deceleration time, and E wave deceleration rate) function before NAC treatment between rats assigned to Lean, HFHS, and HFHS+NAC. Terminal echocardiography and heart dissection revealed group differences in cardiac size and function between groups after ∼8 more weeks of the assigned diet with or without NAC treatment. Cardiac hypertrophy occurred in HFHS rats, but was attenuated in HFHS+NAC rats (Figure 6, Table 2). Percent fractional shortening and posterior wall shortening velocity increased in HFHS relative to Lean controls, and NAC treatment did not lead to a statistically significant change in this phenotype (Figure 6 B-C). E wave deceleration rate (E/DT) was elevated in HFHS rats, while HFHS+NAC did not demonstrate this functional change (Figure 6 D). To explore whether NAC exerted this effect through prevention or treatment of a phenotype, we conducted analyses of pre- and post-NAC treatment pulse wave Doppler images (Figure 7). Notably, HFHS rats significantly worsened in E/DT when allowed to continue on the obesogenic diet without any supplementation (Figure 7 B). There was a trend toward decreased E/DT post-NAC treatment in HFHS+NAC rats (Figure 7 C). The average change in E/DT was statistically different between HFHS and HFHS+NAC groups (Figure 7 D).

**Table 1.** Echocardiographic measurements pre-NAC treatment.

|  | Lean<br>(n = 8) | HFHS<br>(n = 7-8) | HFHS+NAC<br>(n = 8) | ANOVA<br>p-value |
| --- | --- | --- | --- | --- |
| LVIDd (mm) | 8.0 ± 0.8 | 8.4 ± 0.8 | 8.0 ± 0.7 | 0.534 |
| LVIDs (mm) | 4.5 ± 0.8 | 4.4 ± 0.9 | 4.2 ± 0.7 | 0.797 |
| Fractional Shortening (%) | 44 ± 6 | 48 ± 7 | 47 ± 6 | 0.443 |
| PWSV (mm/s) | 40 ± 10 | 40 ± 5 | 39 ± 7 | 0.658 |
| E (cm/s) | 105 ± 17 | 99 ± 21 | 110 ± 12 | 0.470 |
| DT (ms) | 51 ± 10 | 44 ± 9 | 45 ± 8 | 0.218 |
| E/DT (m/s<sup>2</sup>) | 21 ± 3.7 | 23 ± 3.5 | 25 ± 6.3 | 0.280 |
| Heart rate (bpm) | 398 ± 9 | 399 ± 29 | 397 ± 21 | 0.984 |

Data are mean ± SD. P values are for One-Way ANOVA except for posterior wall shortening velocity, which failed normality assumption, so a Kruskal-Wallis One-Way Analysis of Ranks was conducted, and that p-value was reported. LVIDd = left ventricular internal diameter during diastole, LVIDs = left ventricular internal diameter during systole, PWSV = Posterior wall shortening velocity, E = E wave, DT = deceleration time.

**Figure 6.**
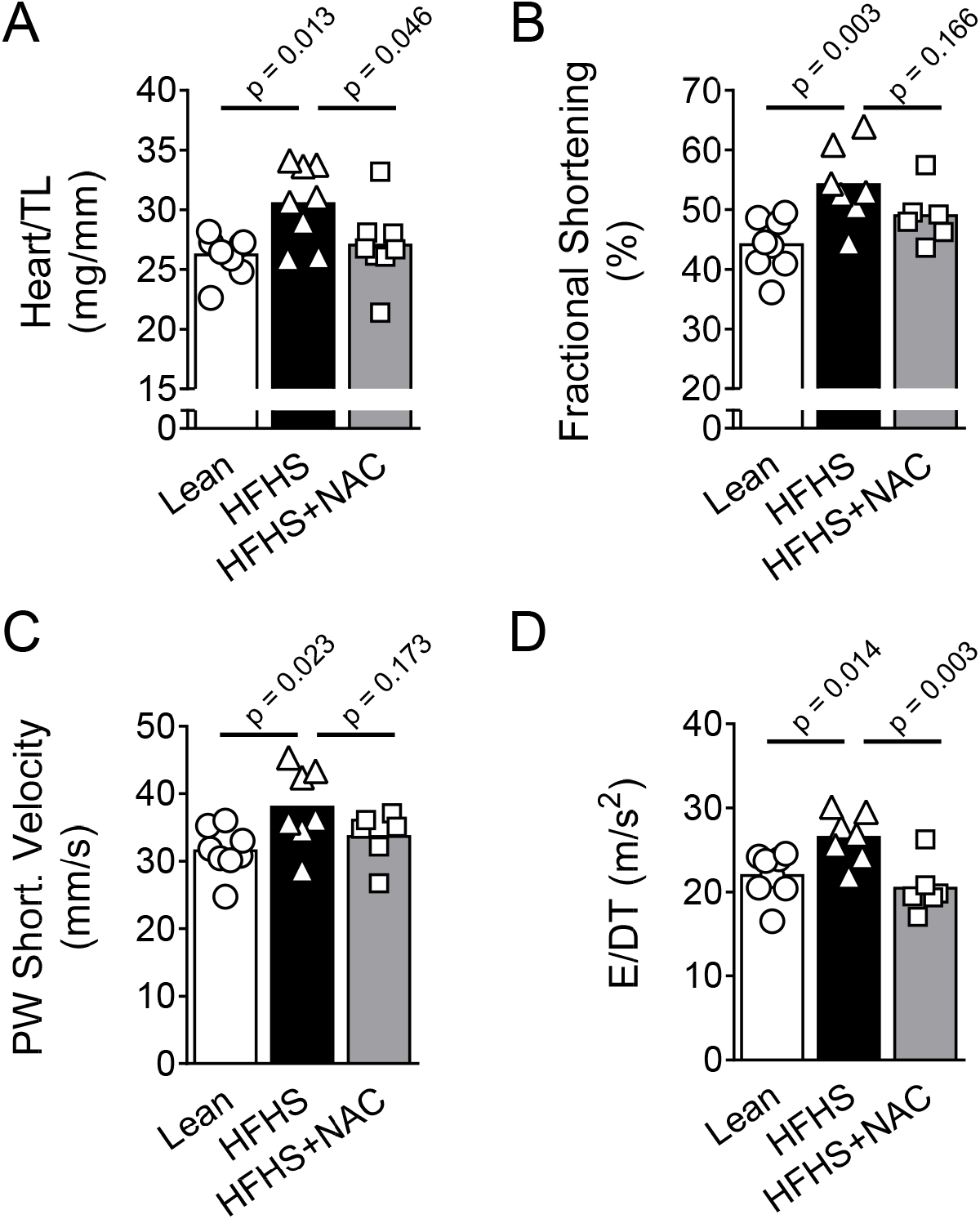
Terminal cardiac size and function. (A) Heart weight normalized to tibia length. Left ventricular (B) fractional shortening (%) and (C) posterior wall shortening velocity. (D) E wave deceleration rate at the level of the mitral valve. Data are shown as scatter plots and mean bars. Comparisons among 3 groups were conducted using One-Way ANOVA. Individual p values for post hoc tests (Dunnett) are shown. TL = tibia length, E = E wave, DT = deceleration time

**Table 2.** Terminal cardiac morphology and function post-NAC treatment.

|  | Lean<br>(n = 8) | HFHS<br>(n = 7-8) | HFHS+NAC<br>(n = 6-8) | ANOVA<br>*p*-value | HFHS vs<br>Lean<br>*p*-value | HFHS vs<br>HFHS+NAC<br>*p*-value |
| --- | --- | --- | --- | --- | --- | --- |
| Heart weight (mg) | 1118 ± 88 | 1314 ± 142 | 1162 ± 131 | 0.011* | 0.008* | 0.040* |
| LV weight (mg) | 894 ± 74 | 1036 ± 118 | 918 ± 99 | 0.020* | 0.017* | 0.049* |
| RV weight (mg) | 224 ± 15 | 278 ± 28 | 243 ± 35 | 0.002* | 0.001* | 0.032* |
| LV weight/TL (mg/mm) | 21 ± 1.5 | 24 ± 2.8 | 21 ± 2.4 | 0.028* | 0.026* | 0.055 |
| RV weight/TL (mg/mm) | 5.3 ± 0.3 | 6.5 ± 0.6 | 5.7 ± 0.9 | 0.004* | 0.002* | 0.041* |
| LVIDd (mm) | 8.0 ± 0.5 | 8.3 ± 0.9 | 7.8 ± 0.4 | 0.559 | - | - |
| LVIDs (mm) | 4.4 ± 0.4 | 3.8 ± 0.9 | 4.0 ± 0.5 | 0.218 | - | - |
| E (cm/s) | 113 ± 7 | 115 ± 24 | 102 ± 18 | 0.361 | - | - |
| DT (ms) | 53 ± 6 | 45 ± 9 | 51 ± 4 | 0.142 | - | - |
| Heart rate (bpm) | 418 ± 21 | 395 ± 24 | 387 ± 24 | 0.050 | - | - |

Data are mean ± SD. P values are for One-Way ANOVA except for DT, which failed normality assumption, and LVIDd and LVIDs, which failed equal variance assumptions, so Kruskal-Wallis One-Way Analyses of Ranks were conducted and those p-values reported. For comparisons that surpassed the threshold for statistical significance ( $p < 0.05$ ), p-values for Dunnett's posthoc tests are also reported. LVIDd = left ventricular internal diameter during diastole, LVIDs = left ventricular internal diameter during systole, E = E wave, DT = deceleration time.

**Figure 7.**
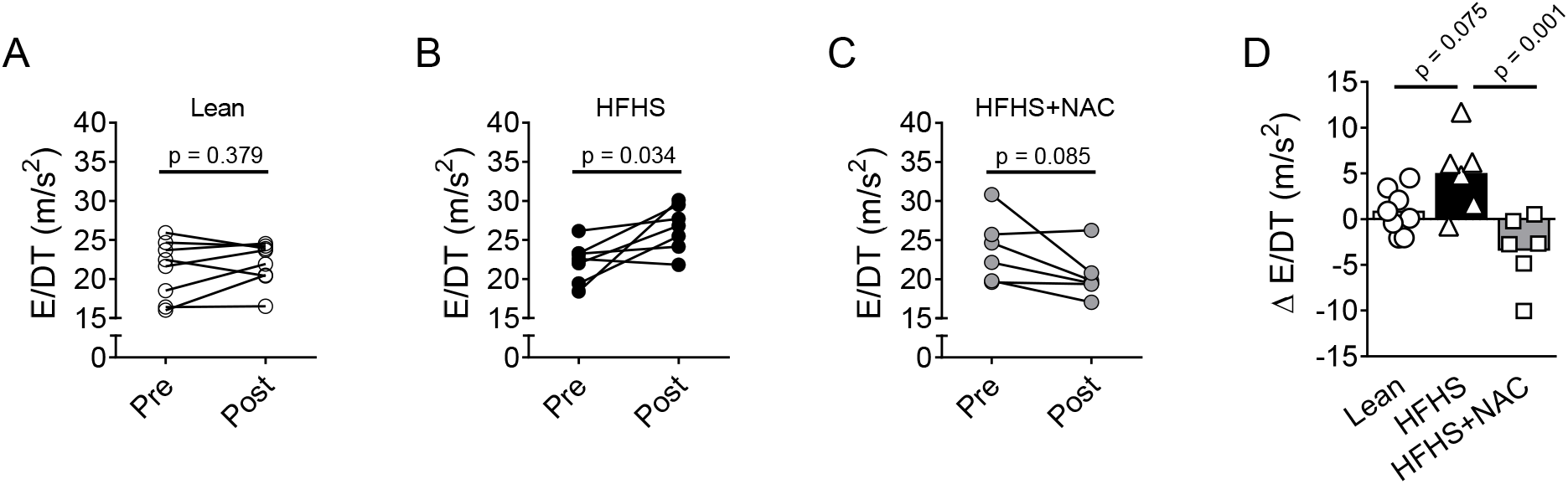
Effects of NAC treatment on diastolic function. E wave deceleration rate (E/DT) at the level of the mitral valve prior to control or NAC-treatment (pre) and after approximately 8-weeks of treatment (post) for lean (A), HFHS (B), and HFHS+NAC (C) rats. Data are shown as scatter plots with lines connecting paired values per animal. Pre and Post values were compared within each group using a paired Student t-test. (D) Change in E/DT shown as scatter plot and mean bars with comparison among the 3 groups conducted using One-Way ANOVA. Individual p values for post hoc tests (Dunnett) are shown. E = E wave, DT = deceleration time

## Discussion

Our main results were that long-term consumption of a high-saturated fat, high-sucrose diet did not result in diaphragm muscle contractile dysfunction or morphological abnormalities but did cause cardiac hypertrophy as well as hypercontractility and diastolic dysfunction. Supplementation with NAC partially attenuated these cardiac abnormalities.

### Diaphragm muscle in obesity and obesogenic diets

There have been previous reports of diaphragm abnormalities in contractile function, morphology, or both in genetically obese rodents, but these data are not consistent across studies. For example, using Zucker fatty rats, Farkas et al.^24^ demonstrated decreased diaphragmatic twitch (∼30%) and maximal specific force (∼13%) and prolonged time to peak tension. On the other hand, using the same genetic obesity model, van Lunteren^64^ showed no change in diaphragm force-frequency relationship or fatigue properties, and De Jong et al.^65^ even demonstrated enhanced diaphragmatic maximal specific force relative to lean controls both before and after mechanical ventilation. In a slightly different strain, the Zucker Diabetic Fatty rat, Allwood et al.^29^ documented respiratory dysfunction as evidenced by lower pressure during maximal occlusion. However, their data showed that this compromise was not associated with changes to diaphragm specific force generation. Additionally, in the *ob/ob* mouse, isolated diaphragm contractile function appears normal.^66^ Nevertheless, some aspect of morphological change has usually been reported in genetically obese rodents, with reports of atrophy,^24,29^ glycolytic to oxidative fiber shift,^24,28^ increased fibrosis,^29^ and/or elevated diaphragm adiposity,^65,66^ but it is notable that the exact pattern of this morphological change varies from study to study.

It is more relevant to compare our diaphragm findings to data from diet-induced obesity studies. Using mice fed a diet with ∼65% of energy from fat, Tallis et al.^25^ observed a ∼30% decrease in maximal diaphragmatic specific force as well as ∼45% compromise in maximal power output. Similarly, Bursas et al.^66^ recently reported diaphragm weakness (∼15% decrease in maximal specific force) when mice were fed the same obesogenic diet as used in our current study (i.e., 45% of energy from fat). These studies also reported morphological changes to diaphragm with diet-induced obesity, showing either a shift toward slow myosin heavy chain isoform expression or increased adiposity and fibrosis.^25,66^ However, the 0.2% increase in polymerized collagen area, which has been reported in diet-induced obese murine diaphragm ^66^, is unlikely to represent a functionally meaningful change.

Discrepant findings of obese diaphragm abnormalities may be related to the severity of metabolic syndrome in rodent models. Our HFHS-fed rats did not exhibit glucose intolerance, and in general, all other previous studies on diaphragm dysfunction have used rodents that exhibited some overt metabolic compromise. The Zucker Fatty rats used by Farkas et al.,^24^ who reported the most contractile dysfunction, were “morbidly obese” and demonstrated a 130% increase in body weight relative to controls. This weight differential far exceeded that of the current study (16%), and even those of other previous studies of rodent diaphragm function in obesity, which have reported body weight differentials of ∼25% to 60% in Zucker rats^28,29,64,65^ or 40% to 50% in diet-induced obese mice relative to controls.^25,66^ Furthermore, several previous studies discussed have demonstrated diabetes or some extent of glucose intolerance with or without hypertension or dyslipidemia.^29,65,66^ Diaphragm isolated from streptozotocin-injected rats, an established model of hyperglycemia, has shown diminished force generation capacity,^67,68^ further suggesting that other aspects of the cardiometabolic syndrome and not weight gain alone are responsible for alterations in the diaphragm. It is possible that if our rats were kept on the HFHS diet longer, they would have developed severe enough cardiometabolic disease that would induce diaphragm dysfunction. This time course issue was demonstrated in mice fed a high-fat diet for 4 weeks that were resistant to limb muscle weakness but, after 12 weeks of feeding, did develop functional and morphological skeletal muscle deficits in the context of hyperglycemia.^69^ We may have failed to reveal any diaphragm differences between Lean and HFHS rats due to Wistar rats’ relative resistance to diet-induced metabolic syndrome, despite their propensity for weight gain. Obesity-prone rat strains and most mouse strains are typically more vulnerable to the cardiovascular, hyperglycemic, hyperinsulinemic, hyperleptinemic, and inflammatory effects of a high-saturated fat, high-sucrose diet,^70,71^ and in mice, this type of diet alone can even promote heart failure.^72^ A complex interplay may exist between the positive training effect of increased body weight on respiratory muscles and the inflammatory milieu of metabolic disease that promotes muscle dysfunction. Therefore, differential severity of the cardiometabolic syndrome may explain some of the equivocal findings across obese diaphragm studies.

Due to our direct, supramaximal stimulation of the diaphragm muscle in the present study, we can conclude that intrinsic muscle dysfunction was not present in our model. However, we did not test for abnormalities upstream of voltage-gated muscle depolarization, such as those in neuromuscular transmission. Neuromuscular transmission failure could explain diminished maximal respiratory pressures in Zucker Diabetic Fatty rats but no difference in specific force values generated by isolated diaphragm bundles. ^29^ Although alteration of the diaphragm neuromuscular junction has not been thoroughly characterized in uncomplicated obesity, data from diabetic rodent models suggest that metabolic syndrome disrupts neuromuscular function and morphology (e.g., acetylcholine receptor distribution).^73,74^ These considerations and our current null results for intrinsic muscle differences support the notion that future studies should evaluate diaphragm function when stimulated through the phrenic nerve.

### Cardiac morphology and function

Our cardiac data recapitulate the most common obesity-related alterations: cardiac hypertrophy and diastolic dysfunction (i.e., elevated E/DT).^33^ Furthermore, we also found evidence of hypercontractility (i.e., higher fractional shortening and posterior wall shortening velocity) with the HFHS diet, which was not evident in the HFHS+NAC group. We believe that NAC likely exerted both ‘preventive’ and ‘treatment’ effects against the cardiac abnormalities reported in this study. HFHS rats had a time-dependent increase in E/DT (worsening of diastolic function) that did not occur in the HFHS+NAC group. Importantlybly, there was a trend toward lower E/DT pre vs. post consistent with a treatment effect.

NAC has direct treatment effects in cardiomyocytes and could also improve cardiac function indirectly. In vitro cardiomyocyte exposure to NAC attenuates hyperglycemia-induced cardiomyocyte toxicity. ^75,76^ In vivo, dietary NAC (4 weeks, same dose as current study) improved maximal force generation and calcium sensitivity of isolated cardiomyocytes as well as an echocardiography marker of diastolic dysfunction (i.e., E/A ratio) in rats with myocardial infarction.^52^ Similarly, NAC treatment in a genetic model of heart failure attenuated cardiac oxidative stress, fibrosis, and wall thickening.^77^ In addition to these effects in cardiomyocytes, dietary NAC can also act through several indirect, non-cardiac mechanisms to exert preventive effects on cardiac morphology and function. Dietary NAC is absorbed in the small intestine and enters portal circulation to the liver, where hepatocytes metabolize it to cysteine and then use it in the synthesis of glutathione.^78^ After this “first-pass” metabolism, the liver releases glutathione into the plasma, where it will be transported to cardiac tissue and other organ systems. There are several examples of NAC improving systemic or non-cardiac physiology that may be relevant to our results. NAC supplementation in drinking water prevented weight gain and hyperglycemia in mice fed the same high-saturated fat, high-sucrose diet used in our study (Research Diets, D12451). ^79^ In a model of diabetic renal disease, NAC supplementation increased renal glutathione, decreased oxidative stress, and improved renal function (i.e., less proteinuria and increased creatinine clearance), independent of changes in glycemic control. ^80^ NAC treatment in a rodent model of binge eating reduced markers of hedonic drive to eat. ^81^ Although NAC did not change caloric intake, a potential normalization of eating patterns would elicit beneficial metabolic and systemic effects. NAC antioxidant actions that lessen ROS-induced microvascular dysfunction and hypertension are also part of potential non-cardiac effects that indirectly ameliorate obesity cardiomyopathy. ^82-85^ Therefore, NAC improvements in other cell types and tissues may have lessened diet-induced pathology in general and contributed to NAC-related prevention of an HFHS cardiomyopathy phenotype.

Obesity leads to activation of the renin-angiotensin-aldosterone system and plasma volume expansion,^86^ which increases the systolic and diastolic measures quantified in this study because of their load-dependent nature.^84^ Indeed, load-dependency is a significant limitation of most echocardiographic analyses.^87-89^ Most echocardiography variables are sensitive to changes in preload (i.e., myocardial stretch), afterload, intrinsic cardiomyocyte dysfunction, or a complex combination of these factors. It is possible then that both the systolic and diastolic functional changes seen in the HFHS rats may be related to the plasma volume increase driven by weight gain and sodium and water retention,^86^ and NAC benefits could be linked to fluid status (i.e., a decrease in plasma volume). Rats in the HFHS+NAC group drank less water than the HFHS group. NAC-treated water has odor and taste that may have discouraged consumption. However, visual and manual inspection suggested that all rats had normal hydration throughout the study. NAC administration lowers urine water excretion in angiotensin II-treated mice,^90^ and direct kidney perfusion with antioxidant promotes water retention in spontaneously hypertensive rats.^91^ The lower water consumption by HFHS+NAC rats may have been a compensatory mechanism to maintain euhydration. In general, more sophisticated measures will be required to define the mechanisms underlying the benefits of NAC supplementation.

### Summary

An HFHS diet that induced moderate weight gain increased heart mass and caused cardiac dysfunction, and NAC partially attenuated these effects. These findings support trials with dietary NAC in individuals with obesity because of the supplement’s low-risk profile and ease of use. We also report the absence of intrinsic diaphragm muscle abnormalities in our clinically relevant HFHS diet. These diaphragm findings, along with previous equivocal data in genetic and diet-induced obesity, suggest that diaphragm-targeted pharmacotherapies or training (e.g., inspiratory muscle resistance) are unlikely the most cost-effective approach to alleviating obesity-related dyspnea. However, inspiratory muscle training may benefit neuromuscular junction problems, which we did not assess in our study.

## Acknowledgments

R.C. Kelley was funded by NIH grant BREATHE T32 HL134621 (principal investigator: Gordon S. Mitchell, University of Florida). L.F. Ferreira was funded by a University of Florida Research Foundation Professorship Award, University of Florida DRPD-ROF2020 grant, and NIH R01-HL130318.

